# Chromosome-scale genome assembly of *Malcolmia littorea* using long-read sequencing and single-pollen genotyping technologies

**DOI:** 10.1101/2025.08.12.669985

**Authors:** Kenta Shirasawa, Kazutoshi Yoshitake, Haruka Kondo, Shinji Kikuchi, Keiichiro Koiwai, Sota Fujii

**Affiliations:** Department of Frontier Research and Development, Kazusa DNA Research Institute, Japan; School of Marine Biosciences, Kitasato University, Japan; Department of Horticulture, Graduate School of Horticulture, Chiba University, Japan; Laboratory of Genome Science, Tokyo University of Marine Science and Technology, Japan; Department of Applied Biological Chemistry, Graduate School of Agricultural and Life Sciences, University of Tokyo, Japan

**Keywords:** Chromosome-scale genome assembly, long-read sequencing, single-pollen genotyping

## Abstract

*Malcolmia littorea*, a member of the family Brassicaceae, is adapted to coastal and sandy environments and has become a model in studies of reproductive barriers. However, genomic resources for the species are limited. Here, with the aim of understanding the molecular mechanisms underlying key traits in *M. littorea*, including its survival under harsh conditions, we present a *de novo* genome assembly consisting of 10 chromosome-scale sequences. We employed a high-fidelity long-read sequencing technology for genome assembly. To anchor the sequences to chromosomes, we developed a single-pollen genotyping method to construct a genetic linkage map based on SNPs derived from transcriptomes of pollen grains, possessing recombinant haploid genomes. We built a genome assembly consisting of 10 chromosome-scale sequences (214 Mb in total) for *M. littorea* containing 30,861 predicted genes. A comparative genome analysis and gene prediction indicated that the genome of *M. littorea* is double the size of the *Arabidopsis thaliana* genome, consistent with a whole-genome duplication followed by gene subfunctionalization and/or neofunctionalization in *M. littorea*. This study provides a basis for research on *M. littorea*, an understudied species with ecological and evolutionary significance.

## Introduction

The genus *Malcolmia* (Brassicaceae), a synonym of *Marcus-kochia*^1^, contains a diverse group of plants, including *Malcolmia littorea,* which is uniquely adapted to coastal and sandy environments. The species is able to thrive in challenging habitats, often characterized by high salinity, nutrient scarcity, and strong winds. However, specialized genetic mechanisms underlying resilience have not been determined in the species. Despite its ecological significance and potential as a model for understanding plant adaptation to extreme environments, only six articles published up to 2025 were retrieved through a search using the terms “*Malcolmia littorea*” and “*Marcus-kochia littorea*” against the National Center for Biotechnology Information PubMed database (https://pubmed.ncbi.nlm.nih.gov), and comprehensive genomic resources are lacking.

In addition to its ecological significance, *M. littorea* has recently been adopted as a model in studies of reproductive barriers. A critical aspect of species delimitation and evolutionary divergence in plants is the presence of reproductive isolation barriers, including interspecific incompatibility. These barriers prevent hybridization between distinct species, thereby maintaining their genetic integrity. In the family Brassicaceae, both self-incompatibility and interspecific incompatibility mechanisms are well-documented, often involving intricate pollen–pistil interactions at the stigma surface, leading to the rejection of "unsuitable" pollen. For instance, studies have identified specific stigmatic proteins, such as Stigmatic Privacy 1 (SPRI1)^2^ and SPRI2^3^, involved in the active rejection of pollen from other species in Brassicaceae. *M. littorea* has been used as the main pollen donor species in these studies; however, comparative genomic analyses are necessary to further elucidate the molecular mechanisms underlying reproductive barriers. Analyses of the genetic basis of interspecific incompatibility are crucial for comprehending evolution and diversification within the family Brassicaceae.

Genome sequencing provides an invaluable tool for dissecting the genetic architecture of adaptive traits, elucidating evolutionary relationships, and guiding conservation strategies.

High-fidelity long-read sequencing, also known as HiFi (PacBio, Menlo Park, CA, USA) sequencing, is one of the most promising technologies to assemble genome sequences at the chromosome-scale telomere-to-telomere level, in which a single contig sequence covers an entire chromosome^4^.

However, in some species with complex genome structures, large genome sizes, high heterozygosity levels, and/or high polyploidy levels, including paleo-ploidy, assembled contig sequences are chunked into short fragments, insufficient to cover a chromosome. To overcome this issue, the Hi–C method has been developed to connect short contigs into chromosome-scale sequences^5^.

Alternatively, genetic mapping, based on classical Mendelian principles, enables anchoring genome sequence fragments to linkage maps to establish chromosome-level sequences^6^. However, genetic mapping requires mapping populations generated by crossing parental lines. Time and space are needed to develop and maintain populations and genetically distinct parental lines. Gametophytes possess haploid genomes containing a single set of chromosomes, which are recombinants of parental chromosomes. Therefore, gametophytes from a single parent can be used as a mapping population. Chromosome-scale genome assemblies have been generated by genetic mapping using gametophytes, including sperms in fish (*Gasterosteus nipponicus*)^7^ and spores in mushroom (*Lentinula edodes*)^8^.

By unraveling the complete genetic blueprint of *M. littorea*, we can gain deeper insights into the genes and pathways responsible for its characteristic stress tolerance and physiological adaptations, its phylogenetic position within the family Brassicaceae, and the molecular underpinnings of its interspecific compatibility or incompatibility with other species. In this study, genome sequencing using a long-read technology, *de novo* assembly via single-pollen genotyping, and comprehensive gene annotation of *M. littorea* were performed, providing a foundational genomic resource for future studies of its biology and evolution.

## Materials and methods

### Plant materials

*M. littorea* (MAL-LIT-1 [Sp-54]) was provided by Tohoku Univ. Brassica Seed Bank (https://sites.google.com/dc.tohoku.ac.jp/pbreed/brassica-seed-bank/brassica-seed-bank). A single plant, Bra27-9, used in previous studies by Fujii et al.^2,3^, was used in this study.

### Chromosome observation

Young flower buds of Bra27-9 were fixed with 1:3 acetate: ethanol for at least 2 days and washed with 70% ethanol. Mitotic chromosome slides were prepared with the fixed buds using the enzymatic maceration–squash method^9^. Chromosome images were captured using an OLYMPUS BX-53 fluorescence microscope equipped with a CoolSNAP MYO CCD camera (PhotoMetrics, Huntington Beach, CA, USA) and processed using MetaVue/MetaMorph v7.8 and Adobe Photoshop CS3 v10.0.1. At least five well-spread chromosome slides were used to determine chromosome numbers.

### Genome size estimation using flow cytometry

Fresh young leaves of approximately 5×D5 mm^2^ in area were chopped with a razor blade for 20 s in a Petri dish containing 400 μL extraction buffer (solution A of the Quantum Stain NA UV 2, Germany) which was added to 1600 μL buffer for DAPI staining (solution B of the Quantum Stain NA UV2, Germany) and then passed through a nylon sieve (40-μm mesh). After 2 min DAPI staining, flow cytometric analysis was conducted using CA II cytometer (Partec, Munster, Germany). *Solanum lycopersicu*m was used as the internal standard throughout the analysis. *Arabidopsis thaliana* ecotype Columbia, a member of the same family (Brassicaceae) as *M. littorea,* was used for genome size comparison. The results were based on the mean of three measurements of material from different individuals.

### Genome sequencing and assembly

Genomic DNA was extracted from young leaves using Genomic Tip (Qiagen, Hilden, Germany). The genomic DNA was sheared to an average fragment size of 30 kb using Megaruptor 2 (Deagenode, Liege, Belgium) in the Large Fragment Hydropore mode. The sheared DNA was used for HiFi SMRTbell library preparation using the SMRTbell Express Template Prep Kit 2.0 (PacBio). The resultant library was separated on BluePippin (Sage Science) to remove short DNA fragments (<15 kb) and sequenced using SMRT Cell 8 M on the Sequel II system (PacBio). The obtained HiFi reads were assembled using Hifiasm^10^ with default parameters. Potential sequences from alternative alleles were removed using purge_dups^11^. Organelle genome sequences, identified by sequence similarity searches of the reported plastid and mitochondrial genome sequences of *Arabidopsis thaliana* using Minimap2^12^, were eliminated. Assembly completeness was assessed with embryophyta_odb10 data using Benchmarking Universal Single-Copy Orthologs (BUSCO)^13^.

### Pollen protoplast preparation

Pollen grain protoplasts were prepared as described in Fan et al.^14^ with minor modifications. Mature pollens were collected from 30 flowers of Bra27-9, washed in pre-buffer solution containing 1/2 MS, 1 M Sorbitol, 0.5 M glucose, and 5 mM MES (pH 5.8), filtered through a nylon mesh (diameter = 70 µm), and suspended in 15 mL of the pre-buffer. After adding 5 mL of 4× enzyme solution (1/2 MS, 1.0% Macerozyme R1-0, 2.0% cellulose Onozuka RS, 1 M Sorbitol, 0.5 M glucose, 0.1% BSA, and 5 mM MES [pH 5.8]) to the pollen suspension, protoplasts were released at 28°C for 3 h at 350 rpm. The mixture containing pollen protoplasts was centrifuged at 100 × *g* for 5 min and pellets were gently resuspended in 10 mL of pre-buffer. This step was repeated twice to remove enzymes and pollen cell wall debris.

### Cell sorting and transcriptome analysis

The Drop-Seq procedure was used to encapsulate single hemocytes and single mRNA capture beads into fL-scale microdroplets, as described previously.^15^ Briefly, a self-built Drop-Seq microfluidic device was prepared by molding polydimethylsiloxane (PDMS; Sylgard 184, Dow Corning Corp., Midland, MI, USA) in the microchannel structure using a negative photoresist (SU-8 3050, Nippon Kayaku Co., Tokyo, Japan). Using this device, droplets containing a pollen protoplast and a Barcoded Bead SeqB (ChemGenes Corporation, Wilmington, MA, USA) were produced up to 2 mL per sample using a pressure pump system (On-chip Droplet generator, On-chip Biotechnologies Co., Ltd., Tokyo, Japan). The number of pollen protoplasts was adjusted to 1.2 × 10^5^ cells/buffer.

Droplets were collected from the channel outlet into a 50 mL conical tube. Then, droplets were broken promptly and barcoded beads with captured transcriptomes were reverse transcribed with 25 µM template switching oligo using Maxima H Minus Reverse Transcriptase (Thermo Fisher Scientific) at 25°C for 30 min, followed by 42°C for 90 min. Then, the beads were treated with Exonuclease I (New England Biolabs, Ipswich, MA, USA) to obtain single-cell transcriptomes attached to microparticles (STAMP). The first-strand cDNAs on beads were amplified using PCR. The beads obtained were distributed throughout PCR tubes (2,000 beads per tube), wherein 1× KAPA HiFi HS Ready Mix (KAPA Biosystems) and 0.8 μM 1st PCR primer were included in a 50 µL reaction volume. PCR amplification was achieved using the following program: initial denaturation at 95°C for 3 min; 4 cycles at 98°C for 20 s, 65°C for 45 s, and 72°C for 6 min; 12 cycles of 98°C for 20 s, 67°C for 20 s, and 72°C for 6 min; and a final extension at 72°C for 5 min. The amplicons were pooled, double-purified with ×0.6 AMPureXP magnetic beads (Beckman Coulter, Brea, CA, USA), and eluted in 35 µL of ddH_2_O. Sequence-ready libraries were prepared using the NexteraXT kit (Illumina, San Diego, CA, USA). A total of 600 pg of each cDNA library was fragmented using a transposome. The amplified library was purified using ×0.6 and ×1.0 AMPureXP beads and sequenced (paired-end) on an Illumina NextSeq 500 sequencer, with 20 cycles for cell barcodes and UMI as read1 with custom sequence primers and 130 cycles for read2.

### SNP genotyping, genetic mapping, and chromosome-level sequence construction

Mapping was performed using the STAR aligner^16^ via the Drop-seq_alignment.sh script from Drop-seq tools v2.5.1 (https://github.com/broadinstitute/Drop-seq). The top 300 cell barcodes with the highest read counts were extracted and saved as individual BAM files. To obtain the ground-truth genotypes, HiFi reads were mapped using minimap2^12^ v2.26 with the -ax map-hifi option, and SNP calling was performed using DeepVariant^17^ v1.4.0 with the --model_type=PACBIO option. The reads from the top 300 cell barcodes were then used for SNP calling with bbmap (https://sourceforge.net/projects/bbmap) through the RNA-seq∼SNPcall-bbmap-callvariants script of PortablePipeline v1.4.0 (https://github.com/c2997108/OpenPortablePipeline), setting -v "ploidy=2 minreads=1". SNPs that were not detected in the HiFi reads were removed as noise. Subsequently, a linkage analysis was performed using the linkage-analysis∼SELDLA script of PortablePipeline with the options -b "-p 0.03 -b 0.03 --NonZeroSampleRate=0.005 --NonZeroPhaseRate=0.1 -r 1000 --RateOfNotNASNP=0.001 --RateOfNotNALD=0.01 --ldseqnum 1 --noNewVcf", enabling chromosome-level genome construction. Finally, manual curation was performed using SELDLA-G v0.9.1 (https://github.com/c2997108/SELDLA-G) to build the final genome sequence.

### Gene and repetitive sequence prediction

Protein-coding genes were predicted using two approaches, an *ab initio* strategy using Helixer^18^ and a homology-based gene prediction strategy using GeMoMa^19^. The gene model of *A. thaliana* Araport11^20^ was employed for homology-based gene prediction. Prediction completeness was assessed with the embryophyta_odb10 data using Benchmarking Universal Single-Copy Orthologs (BUSCO)^13^.

Repetitive sequences in the assembly were identified using RepeatMasker (https://www.repeatmasker.org), based on repeat sequences registered in Repbase and a *de novo* repeat library built using RepeatModeler (https://www.repeatmasker.org).

### Comparative genome structure analysis

Pairwise collinear blocks were detected using MCScanX^21^, with a match score (-k) of 50 and match size (-s) of 50. Results were visualized using a synteny browser, SynVisio^22^.

## Results

### Chromosome number and genome size of M. littorea

A total of 20 chromosomes were observed in somatic cells of *M. littorea* buds (Figure 1), suggesting that the chromosome number of *M. littorea* was 2*n*D=D20. The chromosomes were small and similar in size.

**Figure 1.**
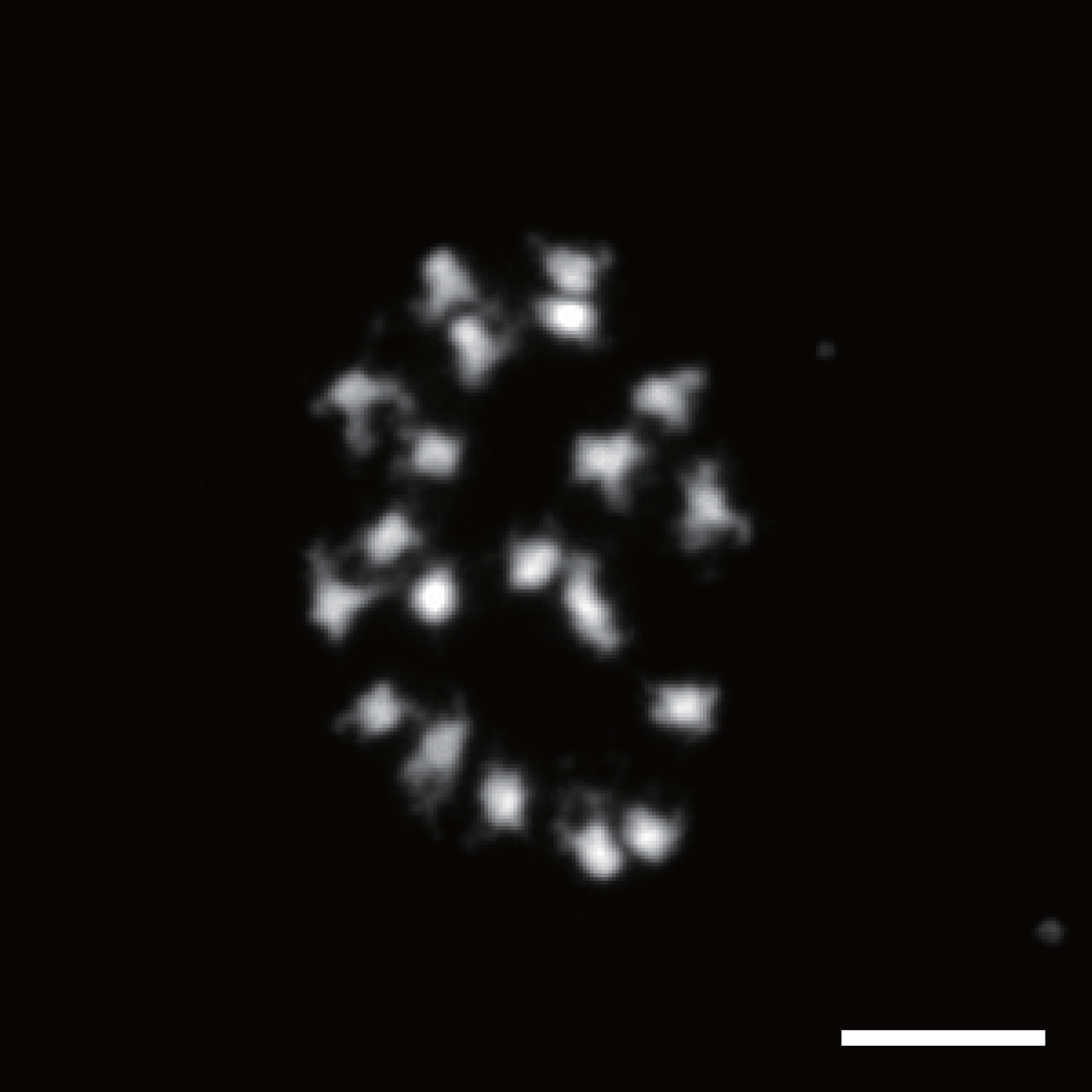
Chromosomes of *Malcolmia littorea*. Chromosomes observed in a somatic cell of *M. littorea* buds. Scale bar = 5 µm.

The genome size of *M. littorea* was estimated by flow cytometry (Figure 2A) using tomato and *Arabidopsis* as controls. The relative genome sizes of *M. littorea*, *Arabidopsis*, and tomato were 0.19, 0.117, and 1.0, respectively. This result indicated that the genome of *M. littorea* was 1.62-fold larger than that of *Arabidopsis* (157 Mb)^23^, corresponding to a size of 254 Mb (= 157 Mb × 1.62).

**Figure 2.**
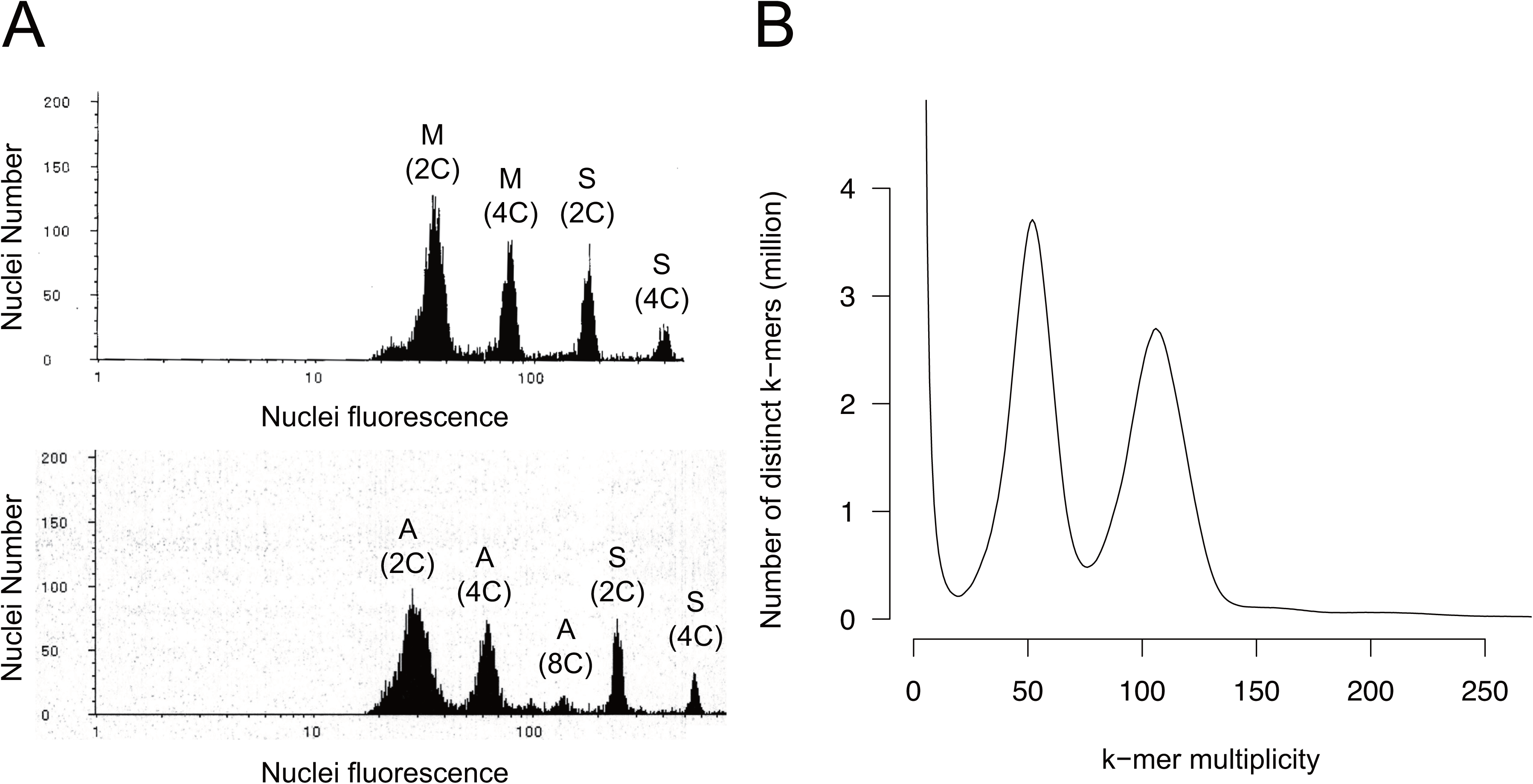
Size estimation of the *Malcolmia littorea* genome. **A.** Flow cytometric analysis of *Arabidopsis thaliana* (A), *Solanum lycopersicum* (S), and *Malcolmia littorea* (M). **B.** *k*-mer analysis (k = 17) with the given multiplicity values.

Furthermore, 26.1 Gb of HiFi reads, with an N50 length of 19 kb, was obtained and used for a *k*-mer distribution analysis (Figure 2B). The *M. littorea* genome was highly heterozygous (1.8%), and the estimated haploid genome size was 246.1DMb. Based on these two analyses, the genome size of *M. littorea* was estimated to be approximately 250 Mb.

### Genome sequencing and assembly

The HiFi reads were assembled into 1,316 primary contigs (N50 = 10.0 Mb) spanning a physical distance of 346.1 Mb. The longer primary contig size than the estimated size and the high heterozygosity of the *M. littorea* genome indicated that the primary contigs were a mixture of two haplotype sequences. Therefore, we purged 1,101 contigs (122.2 Mb) as alternative allele sequences. Potential contaminated sequences from organelles were also removed. The final assembly for the *M. littorea* genome covered 219.7 Mb, consisting of 118 contigs with an N50 of 13.1 Mb (Table 1). The final assembly was designated as MLI_r1.0. The complete BUSCO score for MLI_r1.0 was 98.0%, of which 92.8% were single-copy BUSCOs (Table 2).

**Table 1.** ***Malcolmia littorea* genome assembly statistics**

|  | MLI_r1.0 | MLI_r1.0.pmol |
| --- | --- | --- |
| Total size (bp) | 219,721,536 | 213,500,567 |
| No. of sequences | 118 | 10 |
| N50 length (bp) | 13,098,664 | 23,069,650 |
| N90 length (bp) | 6,584,281 | 12,390,297 |
| Gap length (bp) | 0 | 0 |
| No. of genes | n.a. | 36,880 |

2 n.a.: Not analyzed

**Table 2.** Completeness of the *Malcolmia littorea* genome assembly and predicted genes.

|  | Genome assembly (%) | Predicted genes (%) |
| --- | --- | --- |
| Complete | 98.0 | 97.5 |
| Single-copy complete | 92.8 | 91.9 |
| Duplicated complete | 5.2 | 5.6 |
| Fragmented | 0.4 | 0.2 |
| Missing | 1.6 | 2.3 |

### Chromosome-level sequence construction by genetic linkage of SNPs from single-pollen transcriptome analyses

Protoplasts were prepared from pollen grains harvested from *M. littorea*. Droplets, each of which contained a single protoplast and a Barcoded Bead SeqB (ChemGenes), were generated using a microdevice, as described by Koiwai et al.^24^ Following the protocol of Koiwai et al.^24^, single-cell RNA-Seq libraries were prepared and sequenced to obtain 189.2 million reads.

The sequence reads were split into 3,484,660 subsets in accordance with the barcode sequences and mapped on the MLI_r1.0 with a mapping rate of 63.1%. Among the 3,484,660 subsets, we employed the top 300 subsets (29.9% of all data) for genetic analyses. From the read alignment against MLI_r1.0 as a reference, 330,014 SNPs were detected in the 300 subsets. SNP positions were validated by a mapping analysis of the HiFi reads back onto MLI_r1.0. A linkage analysis of these SNPs resulted in a genetic map with 10 linkage groups, referred to as MLI1ch01–MLI1ch10 (Figure 3) consisting of 41,972 SNPs. The total map length was 744.2 cM (Table 3). Twenty-two MLI_r1.0 contigs (97.2% of the assembly in length) were anchored to the genetic map to establish 10 pseudomolecule sequences as a chromosome-scale genome assembly spanning 213.5 Mb (Tables 1, 3).

**Figure 3.**
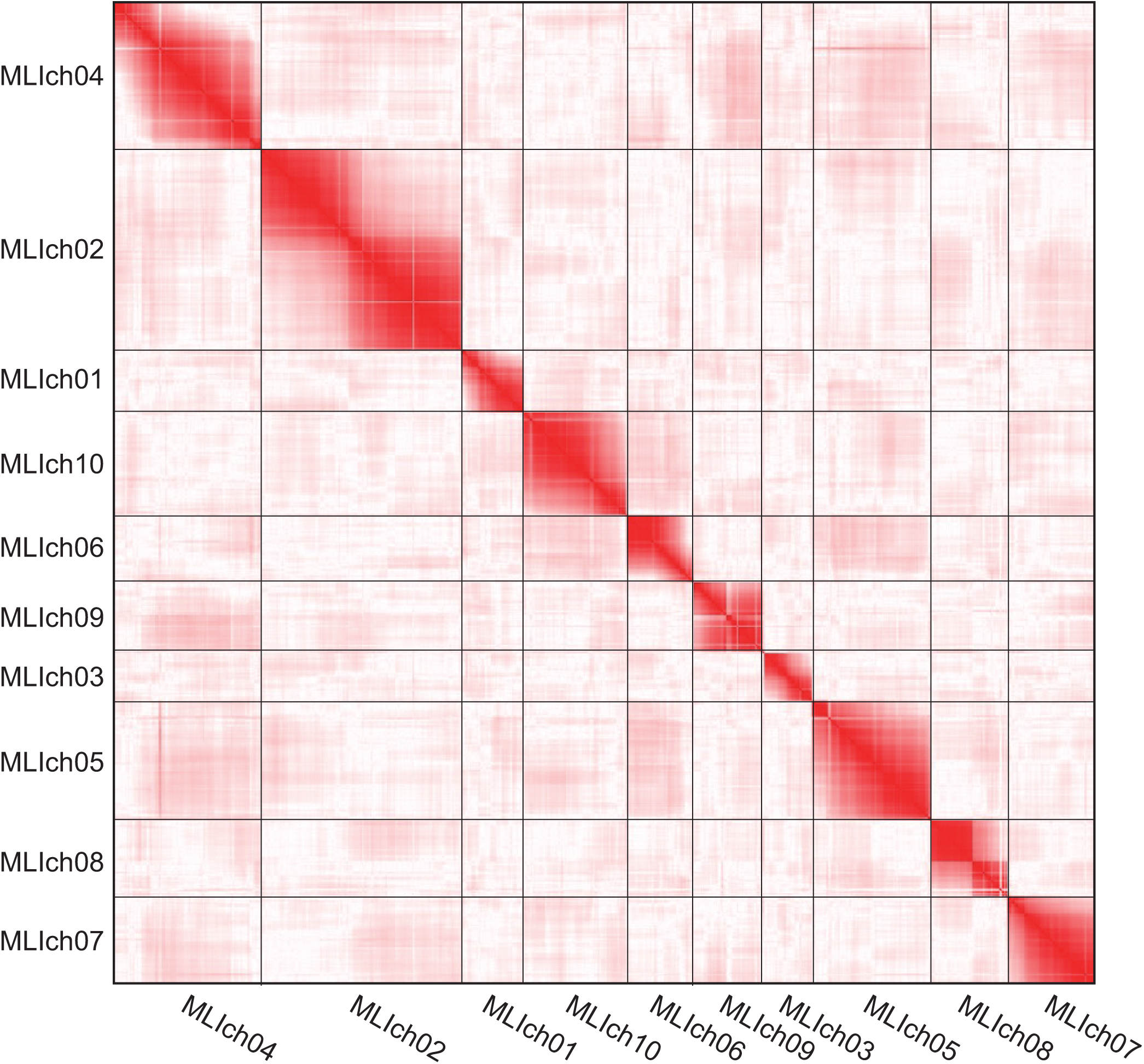
Contact map-style visualization of a single-pollen linkage analysis. Regions in close genetic proximity are shown in red, and distant regions are shown in white.

**Table 3.** **Summary of linkage groups in *Malcolmia littorea***

| Chromosome number | No. of<br>SNPs | Map length<br>(cM) | No. of<br>contigs | Total length (bp) | No. of<br>genes |
| --- | --- | --- | --- | --- | --- |
| MLI1ch01 | 3,289 | 69.9 | 3 | 25,374,001 | 3,305 |
| MLI1ch02 | 7,360 | 72.4 | 2 | 27,430,404 | 4,818 |
| MLI1ch03 | 2,274 | 55.5 | 2 | 20,095,805 | 2,211 |
| MLI1ch04 | 6,536 | 111.6 | 4 | 29,373,502 | 4,280 |
| MLI1ch05 | 4,598 | 119.2 | 3 | 23,069,650 | 3,060 |
| MLI1ch06 | 3,488 | 59.2 | 2 | 21,105,933 | 2,099 |
| MLI1ch07 | 3,462 | 56.8 | 2 | 10,387,743 | 2,276 |
| MLI1ch08 | 3,212 | 49.2 | 1 | 12,390,297 | 2,670 |
| MLI1ch09 | 3,299 | 101.5 | 2 | 20,485,708 | 2,433 |
| MLI1ch10 | 4,454 | 49 | 1 | 23,787,524 | 2,883 |
| Subtotal (ch01 to ch10) | 41,972 | 744.2 | 22 | 213,500,567 | 30,035 |
| Unassigned contigs | n.a. | n.a. | 96 | 6,340,969 | 826 |
| **Total** | **41,972** | **744.2** | **118** | **219,841,536** | **30,861** |

2 n.a.: Not available

### Predicted genes and repetitive sequences in the M. littorea genome

Protein-coding genes in the *M. littorea* genome were predicted through two approaches. First, using the *ab initio* strategy of Helixer, 36,880 gene candidates were obtained. After removing low-confident genes including short exons (1 and 2 bases in length), 26,303 non-redundant genes were selected as candidates. Second, homology-based gene prediction was carried out using GeMoMa, in which 27,655 gene models of Araport11 were employed to obtain 27,803 non-redundant gene candidates. Then, the two lists of gene candidates were merged followed by the removal of redundant genes to obtain 30,861 genes (mean 1,192 bp in length) as a high-confident set (Table 3). The complete BUSCO score for the 30,861 predicted genes was 97.5% (Table 2).

Repetitive sequences occupied a total physical distance of 101.8 Mb (46.3%) in the MLI_r1.0.genome assembly (219.7 Mb). Nine major types of repeats were identified in varying proportions (Table 4). The dominant repeat types in the chromosome sequences were long-terminal repeats (18.0%) including *gypsy*- (13.0%) and *copia*-type retroelements (4.7%). Repeat sequences unavailable in public databases totaled 42.9 Mb.

**Table 4.** Repetitive sequences in the *Malcolmia littorea* genome.

| Repeat type | No. of repeats | Total length (bp) | Relative proportion (%) |
| --- | --- | --- | --- |
| SINEs | 2,715 | 394,972 | 0.2 |
| LINEs | 8,359 | 4,715,176 | 2.1 |
| LTR elements | 42,912 | 39,664,604 | 18.0 |
| DNA transposons | 28,251 | 9,655,138 | 4.4 |
| Small RNA | 3,197 | 716,997 | 0.3 |
| Satellites | 4,149 | 732,066 | 0.3 |
| Simple repeats | 46,962 | 1,899,466 | 0.9 |
| Low complexity | 10,196 | 507,682 | 0.2 |
| Unclassified | 115,777 | 42,933,027 | 19.5 |

### Comparative analysis of the genome structure of M. littorea

Comparative genome analyses identified 92 pairwise collinear blocks between the genomes of *M. littorea* and *A. thaliana* (Supplementary Table S1), ranging from six blocks in MLI1ch03 and MLI1ch06 to 13 blocks in MLI1ch02. Over the genome, two blocks on the *M. littorea* genome corresponded to one block on the *A. thaliana* genome (Figure 4), suggesting a genome-wide duplication in *M. littorea* with respect to the *A. thaliana* genome. Probably due to the genome-wide duplication, 23 pairwise collinear blocks were detected over the *M. littorea* genome (Figure 4, Supplementary Table S2).

**Figure 4.**
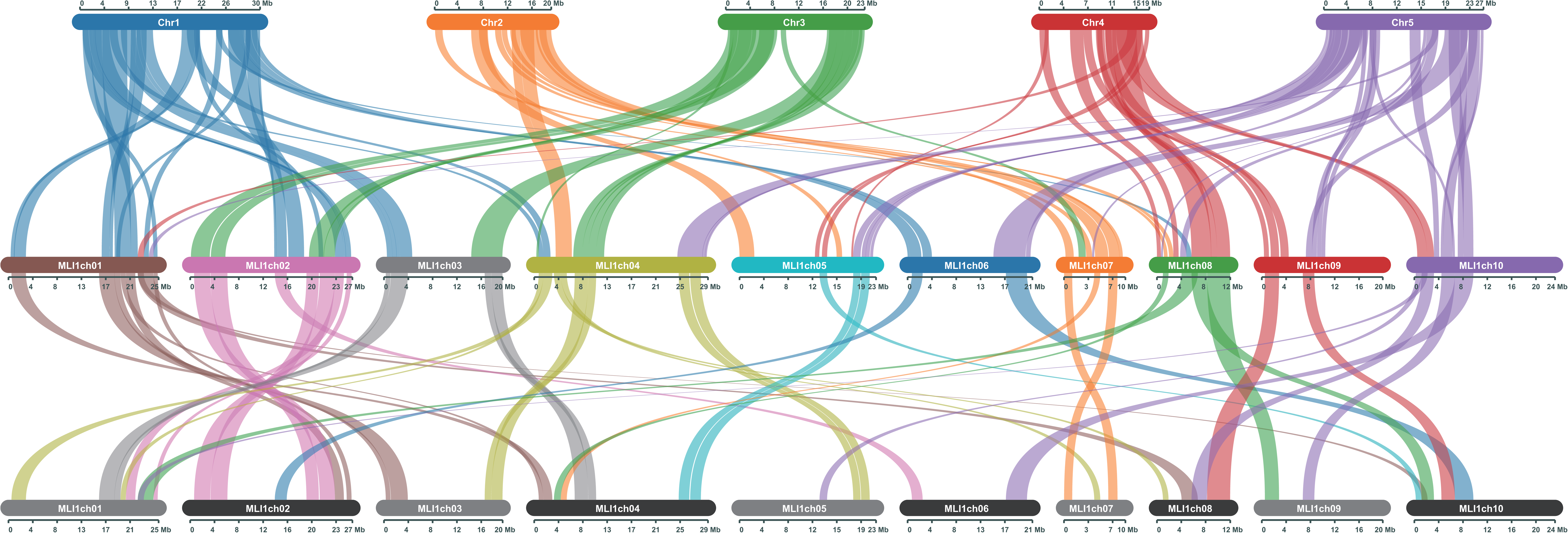
Comparative analysis of genomic structures of *Malcolmia littorea* and *Arabidopsis thaliana*. Chromosome structural similarity between *M. littorea* (MLIch01 to MLIch10) to *A. thaliana* (Chr1 to Chr5) and within *M. littorea*.

## Discussion

Here, we present a genome assembly for *M. littorea* consisting of 10 chromosome-scale sequences, corresponding to the chromosome number estimated through various approaches in this study (Figure 1) and in a previous report^25^. This is the first report of a chromosome-scale genome sequence for *M. littorea* and, more broadly, for the genus *Malcolmia,* except for a draft genome assembly (v1.1) of *M. maritima* registered in a database (Brassicales Map Alignment Project, DOE-JGI, http://bmap.jgi.doe.gov). Comparative analyses of genome structure revealed that the *M. littorea* genome is doubled with respect to the *A. thaliana* genome (Figure 4), even though the genera are closely related^26^. This result suggested that a whole-genome duplication event occurred in *Malcolmia* after its divergence from *Arabidopsis*. Alternatively, *Arabidopsis* underwent rapid diploidization. Interestingly, however, probable duplicated genome segments were highly fragmented and not conserved (Figure 4). Furthermore, the single-copy BUSCO scores for *M. littorea* were 92.8% and 91.9% for the genome and gene sequences, respectively (Table 2), indicating that the gene set in the genome was already diploidized probably via subfunctionalization and/or neofunctionalization. Therefore, *M. littorea* might be a paleopolyploid. These genetic features might explain the ecological adaptation of *Malcolmia* to extreme environments.

For the chromosome-scale assembly, we employed a genetic mapping strategy. Mapping populations are required for genetic linkage analyses. F2 populations can usually be used as mapping populations through self-pollination of an F1 individual derived from a cross between two homozygous lines as parents. F1 or S1 populations generated from crosses between two heterozygous lines or through self-pollination of a single heterozygous line, respectively, can also be used. In this study involving a single line of *M. littorea*, it was impossible to generate such mapping populations due to the self-incompatibility and lack of multiple, genetically diverse lines as parents. To overcome this limitation, we used pollens, each possessing a recombinant haploid chromosome derived from the heterozygous diploid genomes of the parent, as a mapping population.

Single-pollen genotyping has been used in haplotype phasing in genome assemblies in potato^27^ and in pear^28^; however, to the best of our knowledge, this strategy has not been used in genetic mapping, except in fish^7^ and mushrooms^8^. For genome-wide SNP genotyping-by-sequencing, the transcriptome was targeted, even though RNA is transcribed from a small part of genomic DNA (16.8%, calculated from a total gene length of 37.0 Mb divided by the genome length of 219.8 Mb). The abundance of RNA molecules in cells is advantageous over DNAs (only three copies in a single pollen consisting of a vegetative nucleus and two sperm cells) for sequencing. Indeed, we successfully established a genetic map consisting of 41,972 SNPs, which was sufficient to anchor the genome contigs to the genetic map. Therefore, as shown in this study, single-pollen genotyping technology, or single-gametophyte genotyping technology, is useful to establish genetic maps even in organisms with large body sizes, long life cycles, or an inability to be farmed as well as in taxa for which it is difficult to prepare mapping populations.

The chromosome-scale genome assembly generated in this study provides a crucial resource for determining the genetic basis and mechanisms underlying adaptation to extreme environments, interspecific incompatibility, and evolution and diversification within Brassicaceae. Furthermore, this study represents an initial milestone in uncovering the value of neglected species maintained in seed banks.

## Acknowledgements

Seeds of *M. littorea* (MAL-LIT-1 [Sp-54]) were provided by the Tohoku Univ. Brassica Seed Bank. We thank Y. Kishida, C. Minami, K. Ozawa, H. Tsuruoka, and A. Watanabe (Kazusa DNA Research Institute) for their technical assistance.

## Data availability

Raw sequence reads and the assembled sequences were deposited in DDBJ (BioProject accession number PRJDB35718). Gene annotation files for coding sequences, putative peptide sequences, and their genome positions are available at Kazusa Genome Atlas (https://genome.kazusa.or.jp).

